# Changes in the health-related ecosystem services and disservices provided by urban trees over multiple decades in a small city

**DOI:** 10.1101/2025.02.21.639397

**Authors:** Russell Kwong, Thomas E. Victor, Noa Dijstelbloem, Zhou Y. Dong, Ava M. Robillard, Willa A. Gagnon, Christina S. Simon, Camille I. Radhakrishnan, Keily A. Peralta, Elan C. Youngstrom, Emma C. Terjesen, Maia Tsignadze, Daphne Okuyama, Moheng Ma, Daniel S.W. Katz

## Abstract

The effects of urban trees on public health should change over decades due to shifts in tree composition and abundance, environmental conditions, human demography, and disease incidence. However, there are few case studies documenting changes in the health-related ecosystem services and disservices provided by trees over time. Here we use seven tree censuses to quantify changes in the effects of city-owned trees on air pollution, allergenic pollen, hydrology, and heat due to shifts in tree abundance and environmental conditions in Ithaca, New York, a small city in the Northeastern United States. We also review how shifts in disease incidence have affected these trees’ health consequences. From 2005 to 2021, trees removed 20% more ozone and 268% more PM_2.5_; they also removed substantially more runoff and had higher cooling capacity. In contrast, the amount of certain air pollutants removed by trees dropped from 2005 to 2021 for sulfur dioxide (361 kg/yr to 8 kg/yr), carbon monoxide (65 kg/yr to 32 kg/yr), and nitrogen dioxide (380 kg/yr to 298 kg/yr) as their ambient concentrations dropped. Pollen production by street trees from 1947 to 2021 initially dropped due to Dutch elm disease but has quadrupled since the 1980s due to increases in several high-pollen producing genera. Overall, this study illustrates how the public health effects of trees vary over decades due to changes in tree composition and abundance, environmental conditions, and changes in disease incidence and human demography, emphasizing the importance of incorporating long-term perspectives into contemporary tree management decisions.

**Highlights:**

● Urban trees health-related ecosystem services changed over decades in a case study
● The removal of SO_2_, NO_2_, and CO by urban trees is diminishing due to cleaner air
● Shifts in tree composition have had large effects on allergenic pollen production
● Changes in human disease incidence over decades mediate trees health effects

## 1. Introduction

It is well understood that urban trees affect public health (Salmond et al., 2016; Wolf et al., 2020) through ecosystem services such as the removal of air pollutants (Escobedo et al., 2011; Jim & Chen, 2008), heat reduction (Bowler et al., 2010a; Kleerekoper et al., 2012; Moss et al., 2019), flood prevention (Carlyle-Moses et al., 2020; Kuehler et al., 2017), and mental health benefits (Alvarado et al., 2023; Kaplan, 1995). Simultaneously, urban trees also create ecosystem disservices (Roman et al., 2020) that pose public health challenges through the release of allergenic pollen (Cariñanos & Casares-Porcel, 2011; Katz et al., 2024a; Sousa-Silva et al., 2021) and volatile organic compounds (Churkina et al., 2017; Wei et al., 2024) or through falling trees (Conway & Yip, 2016; Way & Balogh, 2022); Table 1. Urban tree management decisions such as tree species selection and planting location therefore have public health consequences that persist through the many decades of a tree’s lifespan. While efforts to maximize the benefits of trees generally focus on current conditions (Nowak & Dwyer, 2007; Roy et al., 2012; Speak et al., 2018), there are expected to be large changes in future environmental conditions (IPCC, 2022), as well as human disease incidence (American Heart Association, 2017; Yaghoubi et al., 2019; Zafari et al., 2021) and demography (Vespa et al., 2020). One way to better understand how the public health effects of urban trees might change in the future is to understand how the health-related ecosystem services and disservices provided by trees have changed over previous decades. However, few such studies exist.

**Table 1.**
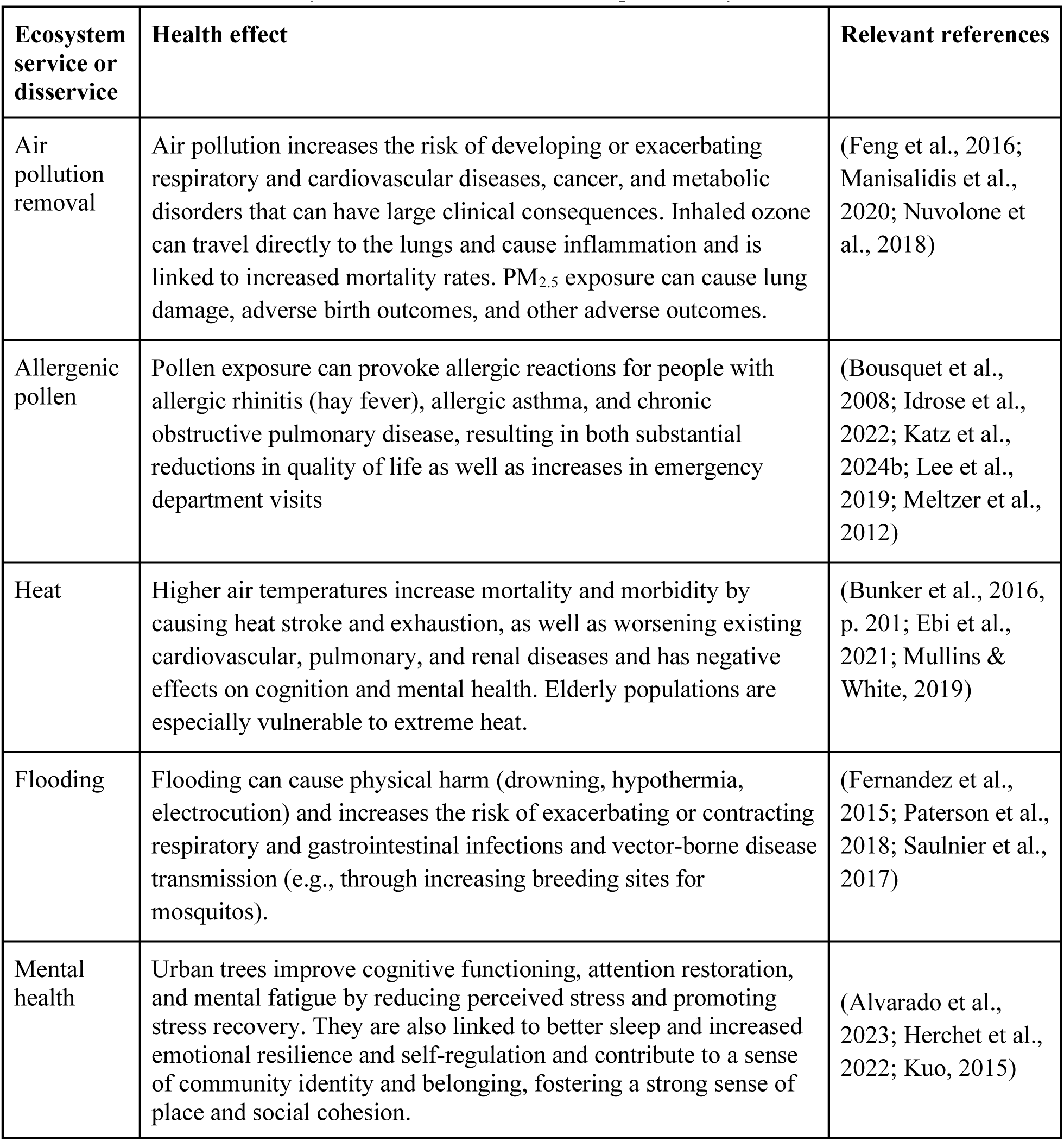
Health effects of ecosystem services and disservices provided by urban trees.

| **Ecosystem service or disservice** | **Health effect** | **Relevant references** |
| --- | --- | --- |
| Air pollution removal | Air pollution increases the risk of developing or exacerbating respiratory and cardiovascular diseases, cancer, and metabolic disorders that can have large clinical consequences. Inhaled ozone can travel directly to the lungs and cause inflammation and is linked to increased mortality rates. PM<sub>2.5</sub> exposure can cause lung damage, adverse birth outcomes, and other adverse outcomes. | (Feng et al., 2016; Manisalidis et al., 2020; Nuvolone et al., 2018) |
| Allergenic pollen | Pollen exposure can provoke allergic reactions for people with allergic rhinitis (hay fever), allergic asthma, and chronic obstructive pulmonary disease, resulting in both substantial reductions in quality of life as well as increases in emergency department visits | (Bousquet et al., 2008; Idrose et al., 2022; Katz et al., 2024b; Lee et al., 2019; Meltzer et al., 2012) |
| Heat | Higher air temperatures increase mortality and morbidity by causing heat stroke and exhaustion, as well as worsening existing cardiovascular, pulmonary, and renal diseases and has negative effects on cognition and mental health. Elderly populations are especially vulnerable to extreme heat. | (Bunker et al., 2016, p. 201; Ebi et al., 2021; Mullins & White, 2019) |
| Flooding | Flooding can cause physical harm (drowning, hypothermia, electrocution) and increases the risk of exacerbating or contracting respiratory and gastrointestinal infections and vector-borne disease transmission (e.g., through increasing breeding sites for mosquitos). | (Fernandez et al., 2015; Paterson et al., 2018; Saulnier et al., 2017) |
| Mental health | Urban trees improve cognitive functioning, attention restoration, and mental fatigue by reducing perceived stress and promoting stress recovery. They are also linked to better sleep and increased emotional resilience and self-regulation and contribute to a sense of community identity and belonging, fostering a strong sense of place and social cohesion. | (Alvarado et al., 2023; Herchet et al., 2022; Kuo, 2015) |

Urban tree composition and abundance shift over decades (Nowak et al., 2016; Roman et al., 2018; Templeton et al., 2019) and each taxon provides a varying mix of ecosystem services and disservices (Masini et al., 2023; Speak et al., 2018). One important agent of urban forest change are exotic pests and pathogens that can extirpate host taxa (Freer-Smith & Webber, 2017; Lovett et al., 2016) and therefore reduce the ecosystem services or disservices these tree taxa provide (Boyd et al., 2013). For example, the emerald ash borer (*Agrilus planipennis*) has decimated ash trees in North America and the associated ecosystem services (Schrader et al., 2021). The changes in urban forests caused by this invasive insect are associated with notable increases in cardiovascular disease and respiratory illnesses (Donovan et al., 2013, 2015). Although less sensational, normal demographic processes such as growth and mortality also alter urban tree populations; individual trees provide varying ecosystem services as they grow, mature, senescence, and die (Katz et al., 2024b; Rötzer et al., 2019).

Many of the public health effects of trees involve regulating the environment, and so the effects of trees also depend on background environmental conditions including air pollution concentrations, temperature, and precipitation. Since the Clean Air Act was passed in 1970, air pollution levels have decreased by 78% in the United States (EPA, 2024a) and air pollution trends continue to change idiosyncratically globally (Sicard et al., 2023). It is possible that the declining air pollution levels in the US over recent decades may have decreased the importance of air pollution mitigation by trees, but these changes have only been investigated at time scales of up to 10 years (Kroeger et al., 2018). Other environmental changes such as warmer temperatures and increases in carbon dioxide concentrations are already increasing the duration of pollen seasons and correlate with increases in allergenic pollen concentrations (Anderegg et al., 2021; Schramm et al., 2021). Climate change is also expected to increase the frequency and intensity of extreme heat events and extreme precipitation events (IPCC, 2022), which could affect the importance of cooling and runoff reduction.

Human disease incidence also shifts on decadal scales (Table 2) and with it, so do the public health consequences of several exposures that trees mediate. For example asthma and other allergic diseases have increased into the early 2000s (Johnson et al., 2021; Pate & Zahran, 2024), due in part to increased urbanization and the adoption of indoor lifestyles (Platts-Mills, 2015). Thus, even if allergenic pollen production did not change over time, its importance would be expected to increase due to societal trends in disease incidence. Similarly, the elderly are especially vulnerable to extreme heat (Bunker et al., 2016; Meade et al., 2020) so an aging population (Vespa et al., 2020) would be expected to increase the importance of the cooling effects of trees.

**Table 2.** Changes in incidence over time in the United States for several diseases that are affected by urban trees.

| Diseases | Relevant (dis)services by trees | Change in incidence over time | Relevant references |
| --- | --- | --- | --- |
| Respiratory disease (asthma, allergic rhinitis, chronic obstructive pulmonary disorder) | Air pollution, pollen | Incidence increased rapidly in the 1960s - 1990s but began to decline again in the 2000s and 2010s; current incidence for asthma is 7.7% | (Binney et al., 2024; Evans et al., 1987; Johnson et al., 2021; Liu et al., 2023; Mannino et al., 1998, 2002; Pate & Zahran, 2024; Platts-Mills, 2015) |
| Cardiovascular disease | Air pollution, extreme heat | Decrease in cardiovascular disease from 1950s through the 2010s; currently responsible for 900,000 deaths per year | (Abdalla et al., 2020; Global Burden of Cardiovascular Diseases Collaboration, 2018; Vasan et al., 2022; Wilmot et al., 2015) |
| Mental health (anxiety, depression) | Various mechanisms | Rapid and ongoing increase starting in the 2000s and 2010s | (Goodwin et al., 2020; Keyes et al., 2019; Weinberger et al., 2018) |

Despite the considerable literature on the effects of urban trees on public health, few studies address how the effects or health-related ecosystem services and disservices provided by trees have changed over time periods of decades. Here, we quantify changes in health-related ecosystem services and disservices provided by trees including air pollution mitigation, allergenic pollen production, cooling capacity, and flood reduction over 77 years in Ithaca, New York, a small city in the northeastern United States. To do so, we calculated the ecosystem services and disservices provided by city-owned trees in seven censuses using i-Tree Eco (Nowak et al., 2008; https://www.itreetools.org), a software suite frequently used to quantify the ecosystem services and disservices provided by trees. We also quantify how the ecosystem services and disservices provided by these trees have changed due to changes in tree composition and abundance and environmental conditions, and provide a brief literature review to contextualize how their health impacts have likely changed due to shifts in human vulnerability (i.e., temporal trends in disease incidence and age). We also describe how trees are expected to become increasingly important for promoting mental health, as anxiety and depression rates surge. Finally, we review how future changes in tree composition, environmental conditions, and human vulnerability are expected to affect the health-related ecosystem services and disservices provided by trees and ultimately how these long-term perspectives should be incorporated into contemporary tree management decisions.

## 2. Methods

### 2.1 Study site description

Ithaca is a small city in the Finger Lakes region of New York State in the Northeastern United States with a population of approximately 32,000 (*U.S. Census Bureau QuickFacts*, n.d.) and an area of 15.7 km^2^. During the academic year, the number of inhabitants approximately doubles due to Cornell University and Ithaca College students (Town of Ithaca, 2024). Ithaca has a humid continental climate (Beck et al., 2023) with an average annual temperature of 7.9°C. The city has warm brief summers, with an average high temperature of 26.6°C, and long cold winters with an average low of −9.7°C. Annual precipitation averages 254.7 cm (Northeast Regional Climate Center, n.d.). Natural plant communities in the Ithaca area are primarily mixed hardwood forests; dominant tree genera include maple, oak, and pine. Ithaca was originally known as “Forest City” and has a long history of publicly supported urban forestry (The City of Ithaca Shade Tree Advisory Committee, 2014).

### 2.2 Tree censuses

Seven city-wide tree censuses have been conducted in Ithaca, in 1928-1947, 1987, 1996, 2002, 2013, 2019, and 2021; these censuses were previously collated and described in detail (Cowett et al., 2021). The 1928-1947 census was conducted sometime in this period by the City Forester; while the exact years of the census are unknown, it likely was conducted over a few years and not over the entire time period (Cowett et al., 2021). For simplicity, we hereafter refer to it as the 1947 census. The 1987 and 1996 censuses were conducted by graduate students at Cornell University under the supervision of city officials, and later censuses were conducted by City Foresters and other city employees. The 1947 census and the 1987 census do not include park trees but these trees were included in all censuses from 1996 onward. Street and park trees from the 1996 census and onwards were distinguished by whether they occurred within City-defined park and natural areas (City of Ithaca, 2024). Although some data are available from 1902, tree size was not recorded so that census is not included in this manuscript. All censuses included here contained, at a minimum, tree species and either diameter at breast height (DBH) or diameter size class. The data from all tree censuses were graciously provided by Cowett et al. (2021) and we used Python and Excel to remove non-tree listings and standardize species names for use with i-Tree Eco (Nowak et al., 2008).

### 2.3 Air pollution calculation with i-Tree Eco

i-Tree Eco was used to estimate the effects of city-owned trees on the removal of criteria air pollutants (PM_2.5_, SO_2_, O_3_, NO_2_, and CO). i-Tree Eco is widely used for this purpose (Riondato et al., 2020; Szkop, 2020; Yao et al., 2022) and uses pollutant specific submodels that have been validated (Hirabayashi et al., 2022); for example PM_2.5_ removal is based on tree leaf area, local wind speed, and current PM_2.5_ concentrations (Kofel et al., 2024; Nowak et al., 2006). The removal of ozone is based on aerodynamic resistance, meteorology, quasi-laminar boundary layer resistance (resistance for ozone entering the air around leaves), and canopy resistance (resistance for air to travel through the tree’s canopy). In our study area, i-Tree Eco could only integrate meteorological and air pollution data from 2005 onwards, as there is not an option to provide custom weather station inputs. Our analysis of the effects of trees on air pollutants and hydrology is therefore restricted to 2005-2021.

Time series of ambient pollution measurements are a key requirement for understanding the effects of trees on air quality, but data were not available for long time periods within our study city. To decide which monitoring station to use in our analyses and how that decision affected our results, we downloaded air pollution data from all nearby air pollution monitoring stations from the EPA Air Quality System API V2 (EPA, 2020) using the ‘RAQSAPI’ R package (Mccrowey et al., 2023) as well as time series for national-scale averages of each criterion pollutant from the EPA (EPA, 2024b). We then selected the nearest station that had both nearly-uninterrupted data and the most similar population density. Specifically, we use PM_2.5_, SO_2_, O_3_, and CO data from Monroe County, New York, which has approximately double the population density to our study area (210 people/km^2^ vs. 86 people/km^2^) and is ∼120 km from our study city; measurements from that location are well studied and representative of cities in the Northeastern US (Emami et al., 2018). For NO_2_, which was not available across that time period from Monroe County, we use data from Lackawanna County, Pennsylvania (population density of 177 people/km^2^), which is ∼130 km from our study city. Although the locations of these air pollution monitoring stations are not from our study city, we display data from the other nearby air pollution monitoring stations to demonstrate consistent regional trends in air pollution concentrations over time. A single outlying measurement from a regional monitoring station (a time series which was not included in the analysis) was omitted from data visualization (NO_2_ in 1996 from Wilkes-Barre, Pennsylvania).

### 2.4 Allergenic Pollen

We calculated pollen production for individual trees as a function of basal area with previously published allometric equations (Katz et al., 2020). We calculated pollen production for *Acer negundo* L., *A. platanoides* L., *A. rubrum* L., *A. saccharinum* L., *Betula papyrifera* Marsh., *Gleditsia triacanthos* L., *Juglans nigra* L., *Morus alba* L., *Platanus × hispanica* Mill. ex Münchh., *Populus deltoides* W. Bartram ex Marshall, *Quercus palustris* Münchh., *Q. rubra* L., and *Ulmus americana* L. Following Katz et al. (2024), species-level equations were applied to congeneric species in the following genera: *Betula*, *Juglans, Morus*, *Platanus, Populus, Quercus,* and *Ulmus*. As per Katz et al. (2024), we did not extrapolate equations for the three *Acer* species in Katz et al. (2020) to the other *Acer* species given the diverse floral morphology within this genus (Rosado et al., 2018). The equation for *Ulmus* used a modified linear equation instead of the exponential listed in Katz et al. (2020) to minimize the effects of outliers. Pollen production was weighted by sex ratios for dioecious taxa, as per sex ratio observations described by Katz et al. (2020).

### 2.5 Cooling

Trees reduce heat through both shading (Rahman et al., 2021; Yu et al., 2020) and transpiration (Winbourne et al., 2020). Trees with more leaves, including both higher leaf area and higher leaf area index (LAI), provide more shade and reduce ground temperature (Speak et al., 2020). Leaf area and LAI are also associated with higher evapotranspiration rates, leading to greater cooling effects (Rahman et al., 2015). Although tree canopy cover has been linked to effects on surface temperature and air temperature at municipal and regional scales (Elmes et al., 2017; Yang et al., 2013), it remains challenging to link individual trees to their total effects on temperature across scales due to the complexities of hydrology, transpiration rates, and atmospheric processes (Li et al., 2024; Massetti et al., 2019) without either extensive ongoing measurements or sophisticated modeling approaches. We therefore use standard allometric equations included in i-Tree Eco to calculate leaf area and LAI for all trees included in the census. These variables serve as rough proxies for the cooling effects of trees; this is a simplification, as it omits the effects of several variables, including transpiration rates, leaf thickness, leaf shape, crown size, and several other relevant variables (Rahman et al., 2020) but is nonetheless more directly associated with cooling potential than generic tree cover, which is used as a proxy for the cooling effects of trees in many larger-scale studies (Y. Yin et al., 2024).

### 2.6 Flooding

Street trees can mitigate the risk of flooding and human health effects caused by floodwaters through capturing precipitation and intercepting groundwater runoff (Aghaloo et al., 2024; Alderman et al., 2012; Livesley et al., 2016). We calculated the hydrology benefits from street trees through i-Tree Eco estimates from 2005-2011. i-Tree Eco has been used extensively to calculate intercepted water and avoided runoff (Hirabayashi, 2013). i-Tree Eco estimates the effects of trees on precipitation interception and runoff with equations that incorporate hourly surface precipitation, evaporation, infiltration, and land cover types. Here we use meteorological data from the nearest local weather station, Ithaca Tompkins International Airport, which is ∼6 km from downtown Ithaca. Data from this station are missing from 2011 to 2013 and are therefore omitted from this analysis. For non-tree census years, we used the nearest future tree census to calculate the hydrology effects of street trees.

### 2.7 Data visualization

We included visualizations of future climate projections for temperature and precipitation for the study area from the Climate Data Explorer (Lipschultz et al., 2020), which includes results from global climate models from CMIP5 that were statistically downscaled with the Localized Constructed Analogs approach (Pierce et al., 2014). Additional statistical analyses and data visualization were completed in R 4.4.1 using several packages within the Tidyverse (Wickham et al., 2019) and in Python 3.13.1. Code for data analysis and visualization are available on GitHub at https://github.com/dankatz/UPPH24.git.

## 3. Results

### 3.1 Changes in tree composition over time

The number of street trees in Ithaca increased from 5,012 in 1947 to 8,354 street trees in the 2021 tree census (Fig. 1A); basal area increased by 25% during that period. Between 1997, the earliest year with park tree data, and 2021, the total number of city-owned trees (street trees and park trees) increased from 9,967 to 11,555; corresponding basal area increased by 46% over that period. In most censuses, the genus with the highest basal area and number of stems was *Acer* (Fig. 1), representing 21.3% of the total street trees in Ithaca in 2021. Within *Acer*, *A. platanoides* was the most abundant and had the largest basal area (SI 1; SI 2). The second most abundant tree genus was *Quercus*, however the genus representing the second largest proportion of total basal area in 2021 was *Platanus*. Likewise, tree genera such as *Salix* and *Tsuga* had fewer but larger individuals.

**Fig. 1.**
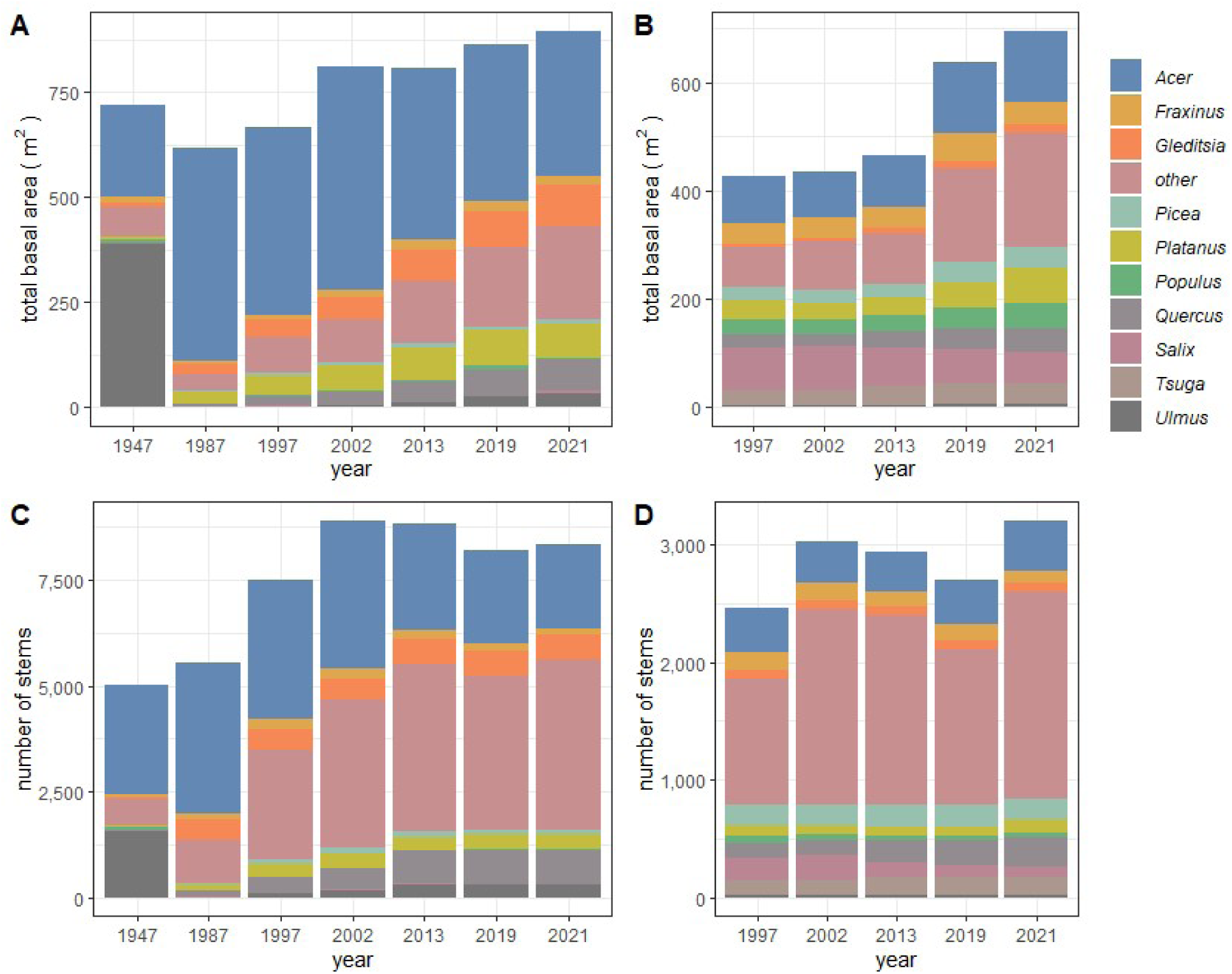
Abundance of the most abundant city-owned tree genera by total basal area for A) street trees and B) park trees, and total number of stems for C) street trees and D) park trees through each tree census.

Changes in street tree composition were also observed over time. For example, *Ulmus* abundance dropped between 1947 and 1987, coinciding with the arrival of Dutch elm disease (*Ophiostoma* spp.).

Recently, ash trees (*Fraxinus*) have started to decline; there were 407 in 2005 and only 255 in 2021. The number of unique street tree genera genus rose from 27 in 1947 to 87 in 2021 (Fig. 1). A full list of species, their abundance, and basal area are provided in SI 3.

### 3.2 Changes in environmental conditions over time

Mean annual maximum temperatures and annual precipitation have varied over the last several decades and temperature is expected to increase in future decades (Fig. 2). Mean annual maximum temperature in the study area is predicted to be approximately 2°C greater in 2060 than in the 1980s (U.S. Federal Government, 2023). Annual precipitation in this area is not forecasted to change considerably but the amount of precipitation is generally expected to become more variable in future climates (IPCC, 2022).

**Fig. 2.**
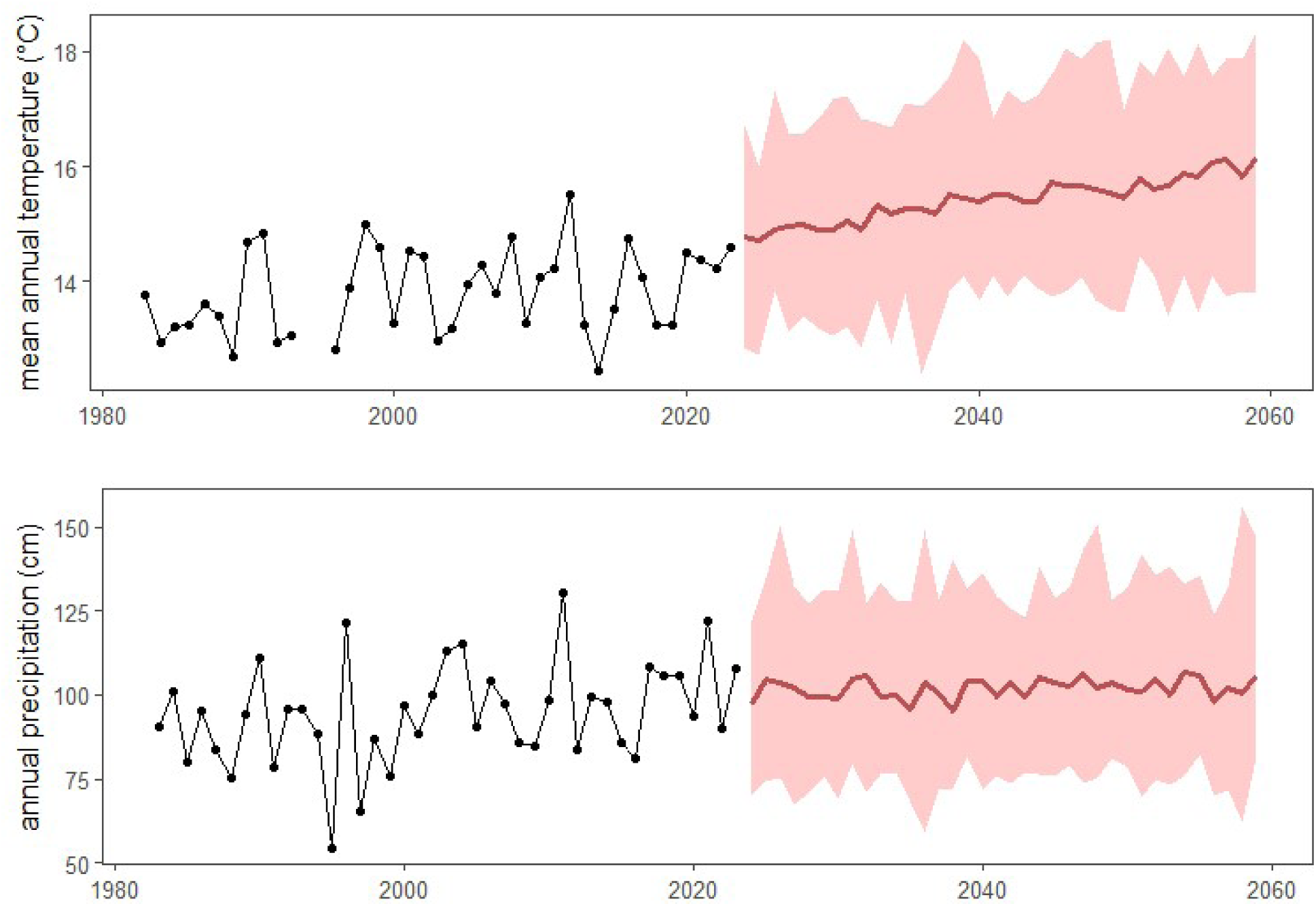
Changes in environmental conditions over time, including historical records from a weather station in Ithaca, NY (black line and dots) for A) annual average maximum daily temperature and B) annual precipitation. Future climate forecasts for the study site are included for the weighted mean of several climate models (red line) and uncertainty (pink shading) based on the RCP4.5 pathway (U.S. Federal Government, 2023).

### 3.3 Air pollution

Criterion air pollution concentrations in the study region have decreased over the last several decades, following national trends (Fig. 3). With the exception of ozone, regional concentrations have generally remained lower than the national average. However, recently, there was a slight rebound in the concentration of PM_2.5_, coincident with the 2023 Canadian wildfires (Byrne et al., 2024).

**Fig. 3.**
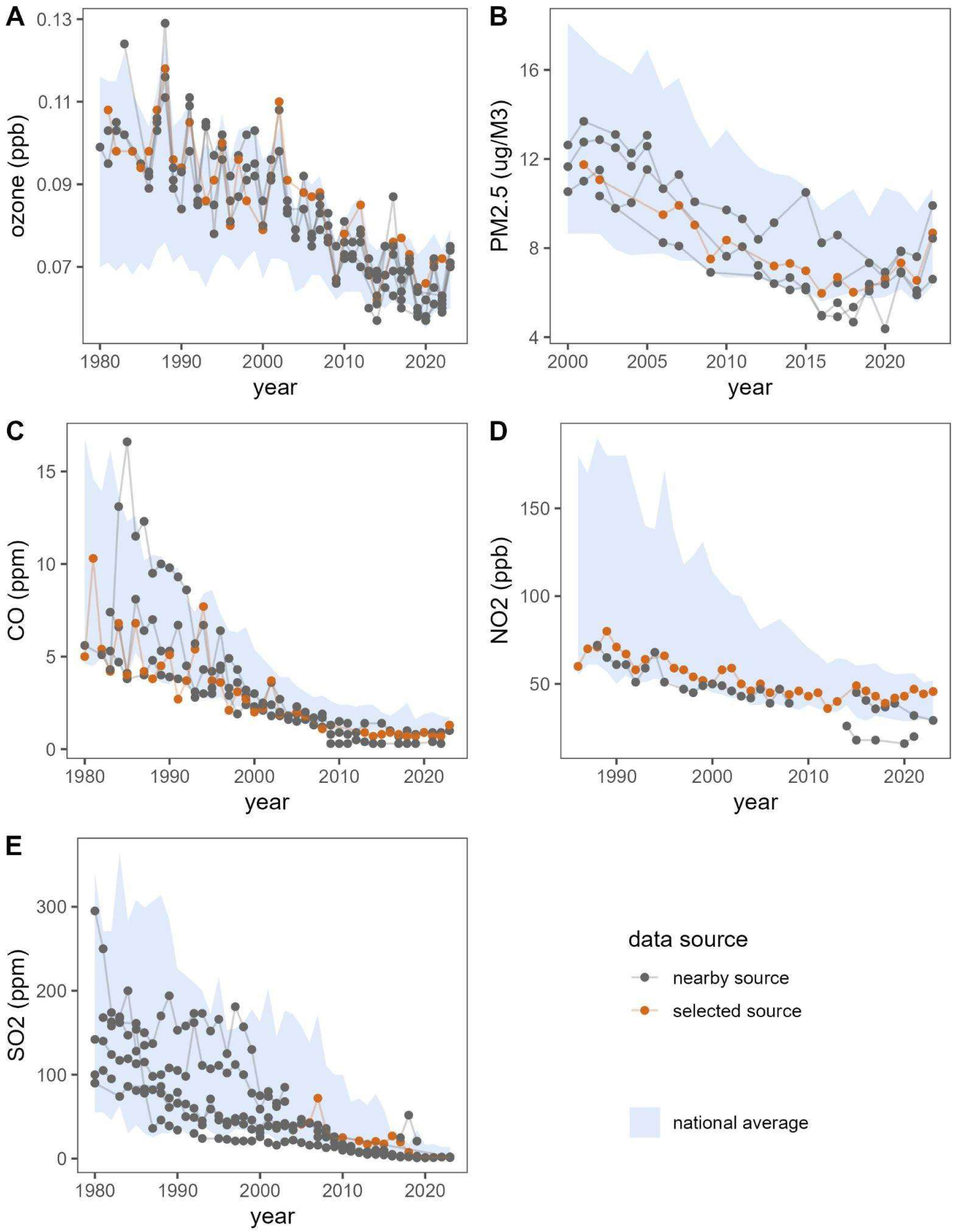
Air pollution concentrations over time at the monitoring sites used in this study (orange), other monitoring sites within the region (gray dots), and the national average (light blue). Air pollution concentrations for years with fewer than 90% completeness are excluded and the specific variables used follow standard reporting nomenclature (O_3_: annual 4th maximum of daily max 8-hour average, PM_2.5_: annual average, CO: annual 2nd maximum 8-hour average, NO_2_: annual 98th percentile of daily max 1-hour average, SO_2_: annual 99th percentile of daily max 1-hour average).

The amount of ozone and PM_2.5_ removed by street trees has steadily increased over the years. From 2005 to 2021, ozone removal rose from 2134 kg/yr to 2551 kg/yr, and PM_2.5_ removal rose from 40 kg/yr to 107 kg/yr, although there has been considerable interannual variation in removal rates. However, the removal of other pollutants has decreased from 2005 to 2021, with sulfur dioxide going from 361 kg/yr to 8 kg/yr, carbon monoxide going from 65 kg/yr to 32 kg/yr, and nitrogen dioxide going from 380 kg/yr to 298 kg/yr (Fig. 4). The total reduction in air pollutant concentrations attributable to the city-owned trees included in the tree censuses in 2005 and 2021 were, respectively, 0.73% and 0.77% for nitrogen dioxide, 1.00% and 1.12% for ozone, −0.02% and 0.38% for PM_2.5_ (negative values are possible from resuspension of PM), 0.97% and 0.54% for sulfur dioxide, and around 0.006% for carbon monoxide in both years.

**Fig 4.**
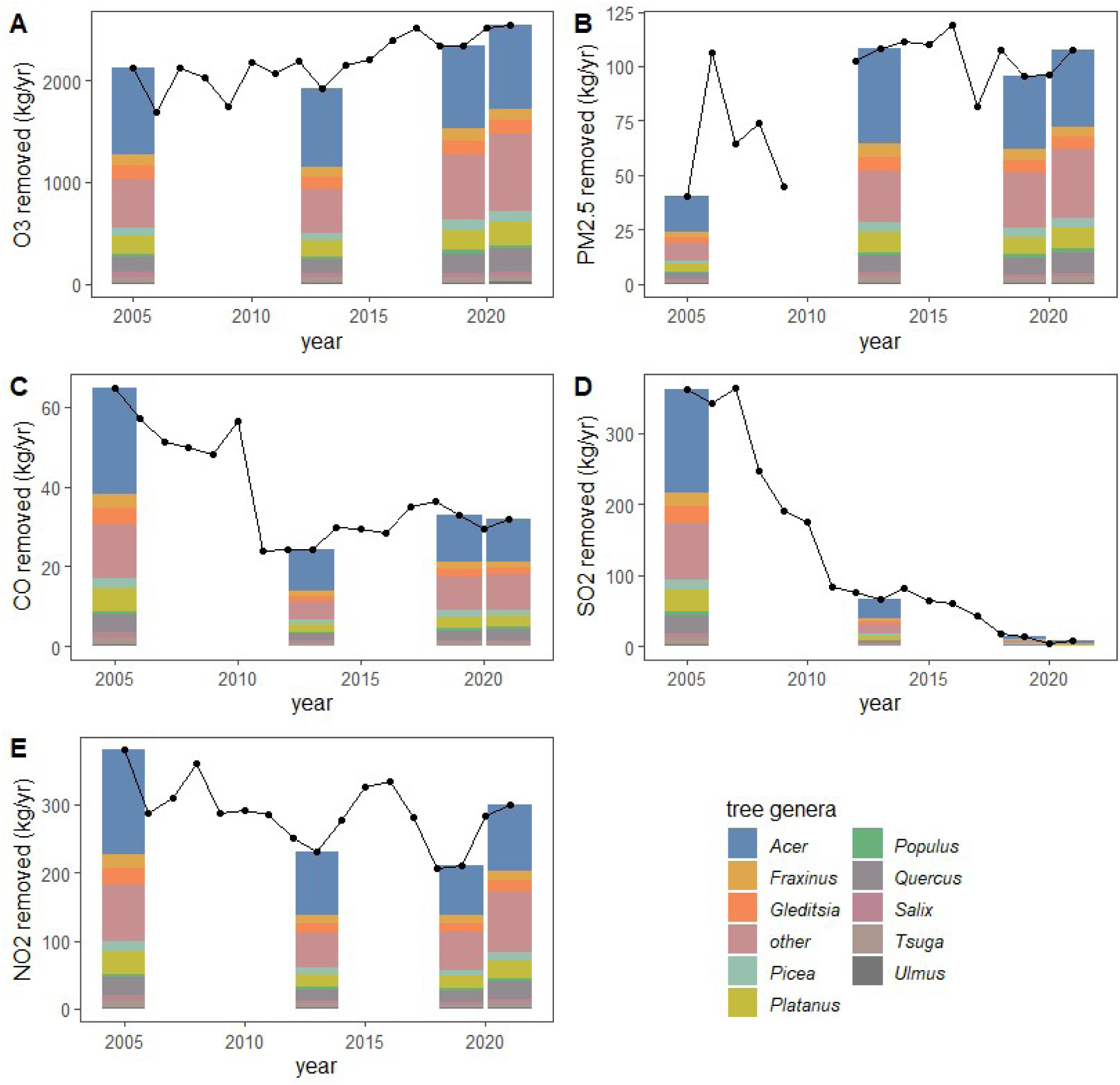
Air pollutant removal by city-owned trees in each census for a) O_3_, b) PM_2.5_, c) CO, d) SO_2_, and e) NO_2_. Removal by each of the top 10 genera by basal area in 2021 are shown; air pollutant removal from other genera are displayed in the ‘other’ category. Air pollutant removal is a function of tree leaf area, ambient air pollution concentrations, and weather conditions. Annual removal rates are shown for years without censuses (black line) using data from the most recent available census. The tree census data from 2002 is shown in 2005, the first year for which i-Tree Eco can be run.

### 3.4 Pollen

Overall, there is an increasing trend in pollen production (Fig. 4) over the last several censuses but there is substantial variation in pollen production among genera. *Platanus* had the highest city-wide pollen production in 2021 (87.6 trillion pollen grains), due to their abundance and large size (Fig. 1) and their high per-plant pollen production. While *Acer* is the most abundant genus, the dominant *Acer* species for which we have allometric equations (*Acer rubrum, Acer platanoides,* and *Acer saccharinum*) generally have low pollen production; *Acer negundo* has much higher pollen production but it is less abundant. Total pollen production for the *Acer* species for which we had allometric equations available was 8.6 trillion pollen grains in 2021. There was a sharp decrease in *Ulmus* pollen production between the first two censuses, which coincides with the arrival of Dutch elm disease in New York.

### 3.5 Cooling

The leaf area of city-owned trees increased by 17.8% from 3.03 million m^2^ in 2005 to 3.57 million m^2^ in 2021. In 2005, the *Acer* genus accounted for 1.21 million m^2^ (40.2%) of total leaf area in Ithaca, and *A. platanoides* was the most prevalent species comprising 15.9% of total leaf area. Following efforts to diversify city-owned trees, *Acer* accounted for 32.3% of leaf area by 2021. While *A. platanoides* remains the most prevalent species, in 2021 it accounted for 11.8% of leaf area. Leaf area of *Gleditsia* and *Quercus* have also increased since 2005. Average LAI values varied considerably by species (SI 4); for example, average LAI for *Acer saccharinum* was 4.6, for *Platanus* x *hispanica* it was 3.3, for *Robinia pseudoacacia* it was 2.4, and for *Malus* species it was 1.7.

### 3.6 Hydrology

In 2021, the amount of runoff avoided and water intercepted have increased by approximately 30% compared to 2005 (Fig. 6). *Acer* consistently reduced runoff by approximately 3,000 m³ per year and intercepted approximately 15,000 m³ of water per year. Species that prevented the largest amounts of runoff included *Acer platanoides* (1,222 m³), *Acer saccharinum* (650 m³), and *Platanus* x *hybrida* (626 m³). Though *A. platanoides* only contributes an annual average of 1.25 m³ of runoff prevented per tree, its prevalence throughout the city accounts for nearly 12% of avoided runoff among all city-owned trees.

**Fig. 5.**
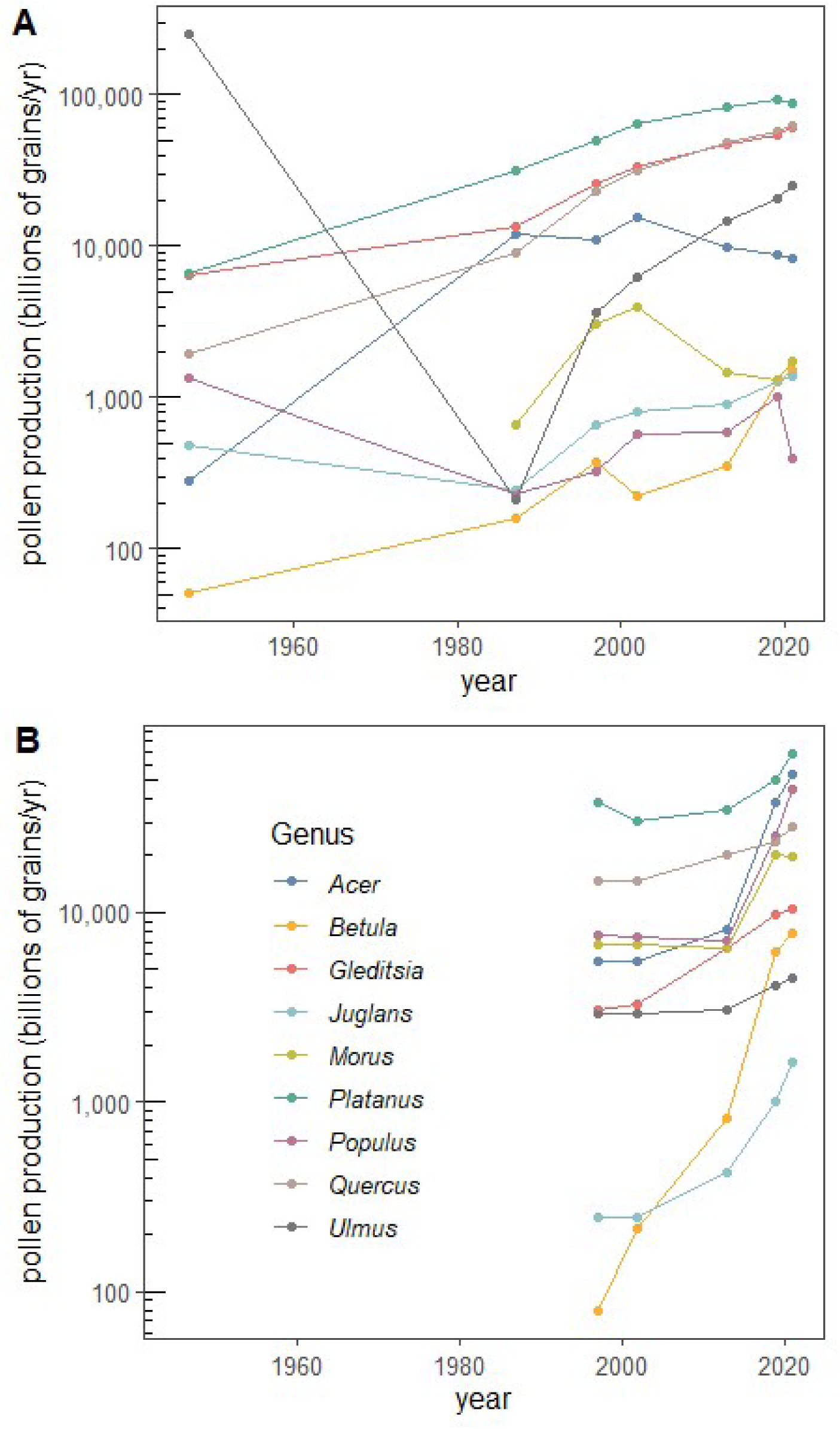
Pollen production (log-scale) by city-owned trees at each census (dots) for the genera that have allometric equations available for A) street trees and B) park trees. Note that pollen production is calculated solely from basal area and does not include interannual variability.

**Fig. 6.**
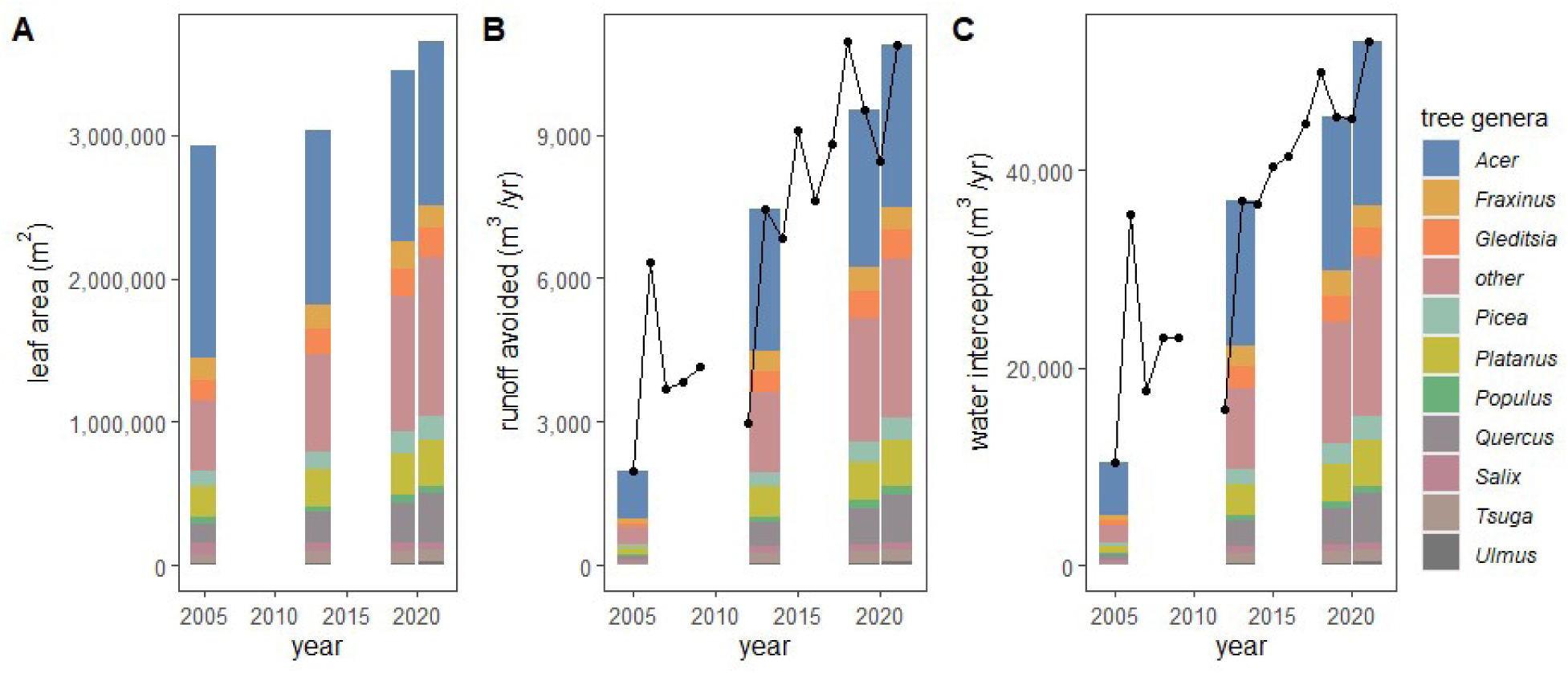
Leaf area and the effects of city-owned trees on hydrology in each tree census for A) leaf area, B) runoff avoided, and C) water intercepted. The effects of each of the top 10 genera (by basal area in 2021) are shown; effects by other genera are displayed in the ‘other’ category. Effects on hydrology are a function of both trees and precipitation and are displayed for all available years (2005-2021). The tree census data from 2002 is shown in 2005, the first year for which i-Tree Eco can be run.

## 4. Discussion

Our results demonstrate considerable changes in the health-related ecosystem services provided in our study area by trees over recent decades. These changes are due to a combination of shifts in city-owned tree composition and abundance as well as concurrent changes in the environment (e.g., reductions in the concentrations of certain air pollutants). Disease incidence and human demography have also shifted over decadal scales (e.g., the rise in allergic disease), which are not included in our quantitative analysis but are reviewed below, as are their anticipated future changes. The health effects of urban trees are the product of multiplicative interactions between trees, the environment, and people, so all three must be traced in tandem to understand changes in the impacts of trees on human health over time. Ultimately, our quantitative results and accompanying review of related literature highlight that optimizing the future health benefits of trees in a changing world will require predictions of future conditions from disciplines including ecology, environmental science, and public health to guide today’s tree planting and management decisions. We also provide a brief review of predictions of future conditions relevant to these ecosystem services and disservices.

### 4.1 Air Pollution

Urban trees in Ithaca removed decreasing quantities of carbon monoxide, sulfur dioxide, and nitrogen dioxide from 2005 to 2021, the period available for modeling in i-Tree Eco. This is largely due to declining ambient concentrations of these gases as legislation such as the Clean Air Act of 1970 and its subsequent amendments have helped reduce emissions and pollution concentrations regionally and nationally (Blanchard et al., 2019; Boubel et al., 1994; Emami et al., 2018; Squizzato et al., 2018). Based on regional air pollution measurements extending back to ∼1980, it can be assumed that trees removed substantially more air pollution prior to 2005. For NO_2_ and CO, these reductions in air pollution concentrations eclipsed the 20% increase in tree leaf area from 2005 to 2021. Changes in the effects of urban trees on O_3_, PM_2.5_, and NO_2_ over this period were moderate and better matched increases in tree leaf area.

While the effects of city-owned trees on air pollution concentrations were very low (ranging between approximately 0 and 1.12% improvement), even small effects on exposures can have noteworthy public health consequences (Nowak et al., 2014), although there is much need for further work on this topic (Eisenman et al., 2019). Despite decreases in asthma and cardiovascular disease incidence over recent decades (Table 2), these diseases remain among the most impactful in the United States, so even small changes in ambient air pollutant concentrations result in large societal health consequences (Ciabattini et al., 2021).

Future changes in the health importance of air pollution removal by trees will depend in part on how policies and other societal changes affect air pollution concentrations. For example, the ongoing shift to electric vehicles could reduce O_3_ concentrations (Schnell et al., 2019) and switching to renewable energy production would further reduce NO_2_ and SO_2_ concentrations (Abel et al., 2018). This suggests that air pollution removal by trees in the United States will continue to decline as will the overall importance of this ecosystem service to human health. That being said, the increasing frequency and severity of wildfires due to climate change is expected to increase the frequency of hazardous smoke exposure events (NOAA National Centers for Environmental Information, 2024). Already, there have been multiple occasions when PM_2.5_ originating from wildfires in Canada has been transported into the study region, raising PM_2.5_ concentrations to extreme levels (Shrestha et al., 2022; Wang et al., 2010; Wu et al., 2018); this is likely to increase in the future (Mansoor et al., 2022). Urban trees also contribute slightly to ozone concentration through biogenic volatile organic compound (BVOC) emissions (Fitzky et al., 2019), which are expected to increase in warmer temperatures (Simon et al., 2019).

### 4.2 Allergenic Pollen

Allergenic pollen production has risen in our study area over recent decades as the number and size of *Platanus*, *Quercus* and several other taxa have increased. This is due in part to the success of Ithaca’s diversification efforts, which have reduced the relative dominance of *Acer*, species of which tend to be relatively low pollen producers. *Ulmus* was the most prolific pollen producer in the first census, but its pollen production dropped precipitously in the 1950s and 1960s when elms in Ithaca were nearly eliminated by Dutch elm disease (Copeland et al., 2022); there were almost no elms in the 1987 census. Since then, *Ulmus* abundance and pollen production have begun to gradually recover but remain an order of magnitude below their peak abundance. *Morus* pollen production was especially variable among censuses, likely because there are few *Morus* trees so total pollen production is more affected by stochastic processes. Although we do not have equations for *Fraxinus* pollen production, it is certainly dropping as the emerald ash borer decimates local *Fraxinus* populations.

Our estimates of pollen production do not account for changes in environmental conditions over the last several decades, which have increased the duration of the spring tree pollen season by days to weeks in North America (Anderegg et al., 2021; Zhang et al., 2015). Analysis of long-term times series of airborne pollen show both positive and negative correlations between average annual airborne pollen concentrations and temperature (Anderegg et al., 2021; Mousavi et al., 2024; Schramm et al., 2021) and some historical correlations from positive correlations have been extrapolated to predict increased pollen production in warmer future climates (Anenberg et al., 2017; Zhang & Steiner, 2022). There is a strong physiological explanation and substantial experimental evidence that elevated carbon dioxide concentrations are increasing reproductive productivity of trees, including pollen production (D’Amato et al., 2007; Kim et al., 2018; Ladeau & Clark, 2006). These environmental changes have presumably already resulted in the production of more pollen by trees globally, including in Ithaca; this trajectory is expected to continue into the future.

The incidence of asthma and allergies rose in the 1950s through the 1990s in the US, although rates have been fairly stable or have declined slightly in the 2000s and 2010s (Binney et al., 2024; Evans et al., 1987; Johnson et al., 2021; Liu et al., 2023; Mannino et al., 1998, 2002; Pate & Zahran, 2024; Platts-Mills, 2015); Table 2. Thus, we expect pollen exposure to have been a trivial ecosystem disservice until the last few decades; for example, there were unlikely to have been many people whose allergies would have benefited from a decrease in *Ulmus* pollen. However, allergenic pollen is now an ecosystem disservice of substantial concern, especially as it becomes easier to link individual trees to the symptoms experienced by individual people (Katz et al., 2023). Increased attention to this ecosystem disservice by the public and urban foresters could reduce the planting of high-pollen species and cultivars (Cariñanos & Casares-Porcel, 2011; Green et al., 2018). Regardless, the planting decisions made in previous decades and those made today will continue to affect local allergenic pollen exposure for many decades to come (Katz et al., 2024a)

### 4.3 Cooling

Although we do not have direct measurements on the cooling effects of trees in Ithaca, we do have information on their leaf area and LAI, which are correlated with cooling capacity (Rahman et al., 2021). From 2005 to 2021, leaf area increased by 17.8%, with presumably proportional effects in tree cooling capacity. Over time, there have also been shifts in species abundance (SI 2) and some of the changes are relevant to cooling, such as the decrease in *Acer saccharum* trees, which tended to have higher average LAI (4.2), or the increases in *Gleditsia triacanthos* trees that had relatively low average LAI (2.8). Based on the current trajectories, it seems likely that Ithaca will see continuing increases in tree leaf area and therefore cooling capacity even as the health impacts of this ecosystem service are growing.

The cooling effects of trees will become more important as extreme heat events become more frequent and intense. In Tompkins County, there has been an increase in days per year with maximum temperatures above 32° C from 1979 to 2016, with higher than usual temperatures in the summertime from the 30-year norm baseline of 24° C (New York State Department of Health, Center for Environmental Health, 2019). Climate change is projected to increase average air temperatures and the frequency of heat waves globally (Rendon et al., 2024) and locally, by 2060, mean annual temperatures are expected to be 2°C greater than 1980 temperatures (Fig. 1).

Nationally, deaths attributed to extreme heat have increased from 495 in 2000 to 2,325 in 2023 (Howard et al., 2024). Locally, there has been an increase in the number of heat-related illness hospitalizations and ED visits in Ithaca, growing from an average of 11 visits over the summer months of May through September in 2008, to an average of 17.7 visits in 2016 (New York State Department of Health, Center for Environmental Health, 2019). Extreme heat is also a growing health risk due to a shift towards an older population nationally (Vespa et al., 2020) and therefore a more vulnerable population. While the increasing prevalence of air conditioning has somewhat reduced heat-related mortality (Sera et al., 2020), concerns remain about co-occurrence of extreme heat events and power outages (Do et al., 2023), which are also likely to increase due to climate change and an aging electrical grid. In urban areas in the United States, there are associations between race, poverty, low tree cover (due in part to structural racism e.g., redlining), and heat (Locke et al., 2021; McDonald et al., 2024), implying that the maximum health effects of cooling by trees could be realized by increasing tree cover in the neighborhoods with the fewest trees and highest heat vulnerability.

### 4.4 Hydrology

Urban trees help mitigate the effects of flooding through intercepting precipitation and reducing runoff (Livesley et al., 2016; Pearlmutter et al., 2017) and the effects of city-owned trees on local hydrology has increased by approximately a third from 2005 to 2020. This outpaces increases in both basal area and leaf area, and is in part due to increases in species that are especially effective at intercepting and capturing water. For example, *Platanus x hybrida* has become more important due its high amount of runoff diversion and increased abundance in recent years.

While climate change is not expected to substantially alter mean annual precipitation in the study area, it is expected to increase variability in precipitation, which contributes to flooding. Indeed, precipitation from extremely heavy storms has increased 70% in the last 70 years (EPA, 2016). As floods are becoming more common with climate change, urban tree management can be a straightforward and economic approach to alleviating health hazards posed by flooding (Zabret & Šraj, 2015). Simply planting trees is recognized as an inefficient blanket solution to flood mitigation but when conducted in combination with other nature-based solutions such as increasing pervious land cover, tree planting can contribute to future flood regulation (Marapara et al., 2021; Wübbelmann et al., 2023).

### 4.5 Mental Health

There is strong evidence that trees are both associated with and improve mental health outcomes (Alvarado et al., 2023; Herchet et al., 2022; Kuo, 2015), but we were unable to quantitatively extrapolate these relationships across regions and cultures to our study area. Nonetheless, the balance of evidence suggests that ongoing increases in city-owned tree abundance is increasing the potential for trees to improve residents’ mental health. Moreover, street tree diversity is correlated with improved mental health outcomes (Wood et al., 2018); in Ithaca, street tree diversity has also increased substantially over the study period (Cowett et al., 2021).

Mental health has decreased across the United States over the past decades (Theriault et al., 2020) and presumably locally too. In Tompkins County, the county containing Ithaca, 21% of adults reported having been told by a health care professional that they have had a depressive disorder and 41% of teenagers in Tompkins County reported that they had felt depressed or sad most days in 2018 (Tompkins County Whole Health & Cayuga Medical Center, a member of Cayuga Health, 2022). Loneliness is a growing epidemic in the US that leads to greater rates of anxiety, depression, and dementia (Office of the Surgeon General, 2023) and trees and tree stewardship can increase social cohesion (Maas et al. 2009; Elmendorf, 2008; Liu, et al 2020; Heid, et al. 2024). The increasing incidence of mental health disorders suggests that the potential mental health benefits of trees have risen substantially in recent years and, based on current trajectories, will be even more important in the future. However, it remains challenging to link particular plants to their mental health effects, and therefore difficult to consider these decisions in tree management and planting decisions.

### 4.6 Limitations

While i-Tree Eco is one of the most commonly used programs for estimating the impact of trees on various ecosystem services and disservices, it has several limitations including its non-spatial framework. It is well understood that air pollution and pollen concentrations vary considerably over small spatial scales (e.g., tens of meters to kilometers; Apte & Manchanda, 2024; Katz et al., 2023). These local gradients are especially pronounced in urban areas where emissions are spatially variable due to emission patterns (e.g., from traffic for air pollution) and air flow patterns (Di Sabatino et al., 2018). In general, the ecosystem services and disservices provided by trees are not distributed evenly and we would not expect vulnerability or the ultimate health effects of trees to be either; this remains an area of ongoing research.

Spatial heterogeneity in air pollution concentrations means that the location of monitoring stations even within a city can impact the estimate for total removal of each pollutant (Pace et al., 2021; Szkop, 2020); here, this is undoubtedly exacerbated by our use of non-local stations. However, trends in air pollution concentrations over time were regionally consistent, suggesting our broader findings of the effects of trees on air pollution over time are robust, even if there is less certainty about the absolute amounts of pollutants removed.

In this study we only focus on city-owned trees, which often comprise approximately 10% of urban trees (McPherson et al., 1997, 2016) and presumably somewhat similar proportions of ecosystem services and disservices. While this does mean that our data only comprise a small portion of all trees in the study area and therefore only a portion of the ecosystem services and disservices provided by all trees in Ithaca, city-owned trees are centrally managed and can most directly incorporate new information directly into policies and management strategies.

## 5. Conclusion

There have been substantial changes in the composition and abundance of city-owned trees over 77 years, including increases in tree abundance and diversity (Cowett et al. 2021) and the near-elimination of certain tree taxa by invasive pests or pathogens. Overall, increasing tree abundance resulted in gains in health-related ecosystems services (removal of O_3_ and PM_2.5_, cooling potential, and reductions in surface water runoff) and in allergenic pollen production. These changes were mostly the result of intuitive tree demographic processes such as planting, growth, and survival that can be readily predicted by urban foresters. However, there were also interactions with environmental conditions beyond the domain of urban forestry, such as the reduction in ambient concentrations of CO, NO_2_, and SO_2_, which resulted in trees removing lower absolute quantities of these air pollutants over time. Similarly, the likely health effects of these ecosystem services and disservices have changed over the decades as human disease incidence and demography have shifted (e.g., the rise of allergic disease and allergenic pollen production or recent declines in mental health). These trends in human vulnerability, while not directly measured in the present study, are well described at national scales. Our results therefore highlight that changes in the public health consequences of urban trees depend not just on the trees themselves but interact with the environment and human disease incidence, which are also in flux. Thus, when making decisions with long-term ramifications, such as tree planting, efforts to maximize the public health benefits of trees should also draw upon cross-disciplinary predictions about future changes in the environment and society.

## Acknowledgements

We are grateful to Fred Cowett, Nina Bassuk, Jeanne Grace, and Kevin Vorstadt for graciously providing us with historical tree surveys in Ithaca, as well as Fred’s insightful comments on an earlier version of this manuscript. We also thank David Miller and Hannah Zonnevylle for additional suggestions. This manuscript is a product of the Urban Plants and Public Health class (PLSCI 4450/6450) at Cornell University.

## Supporting Information

**Supporting Information 1:**
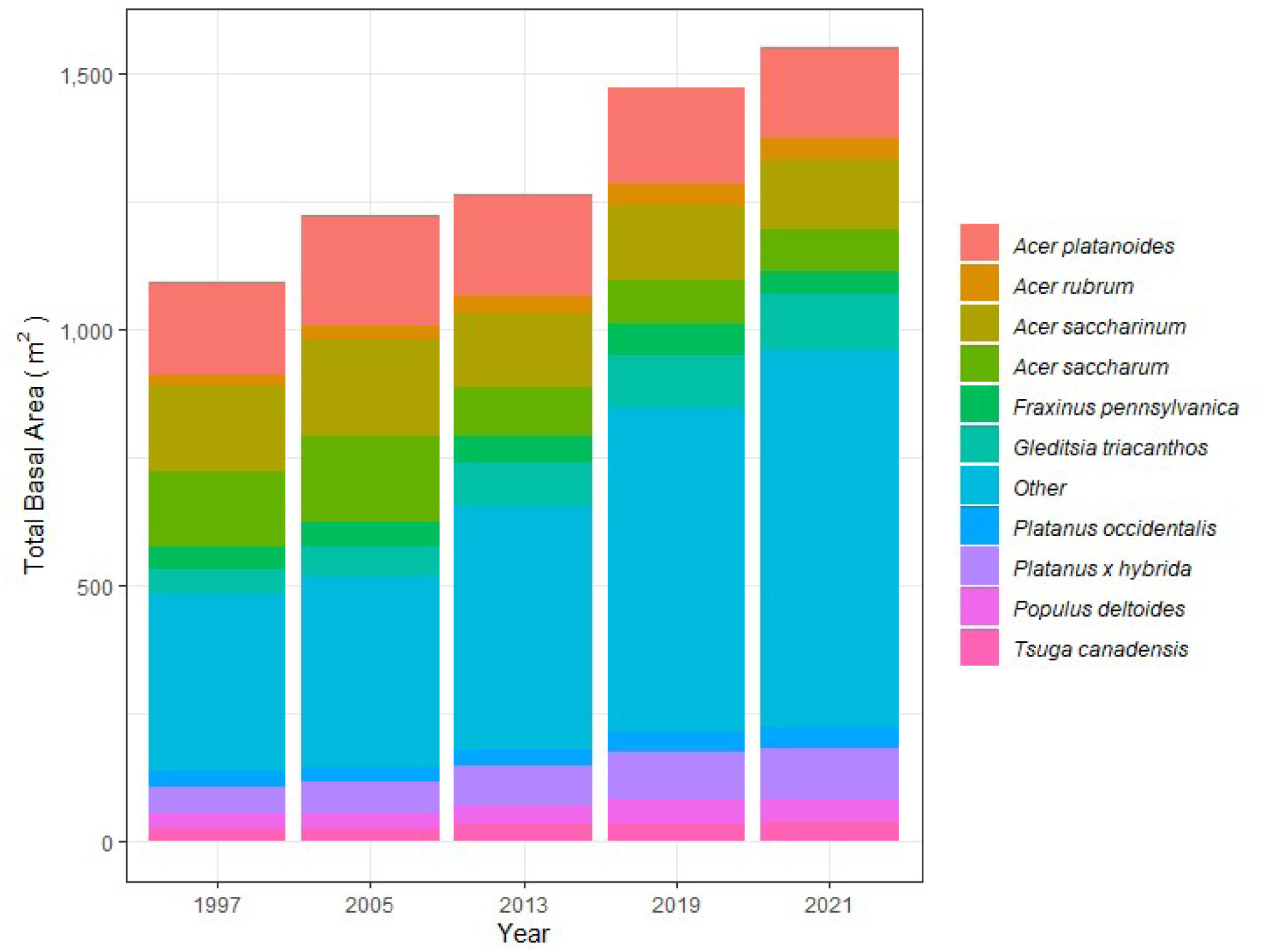
The total basal areas of street trees in Ithaca, New York by species throughout the most recent five tree censuses. The 1928-1947 tree census was omitted as most of the trees recorded were only identified to genus.

**Supporting Information 2:**
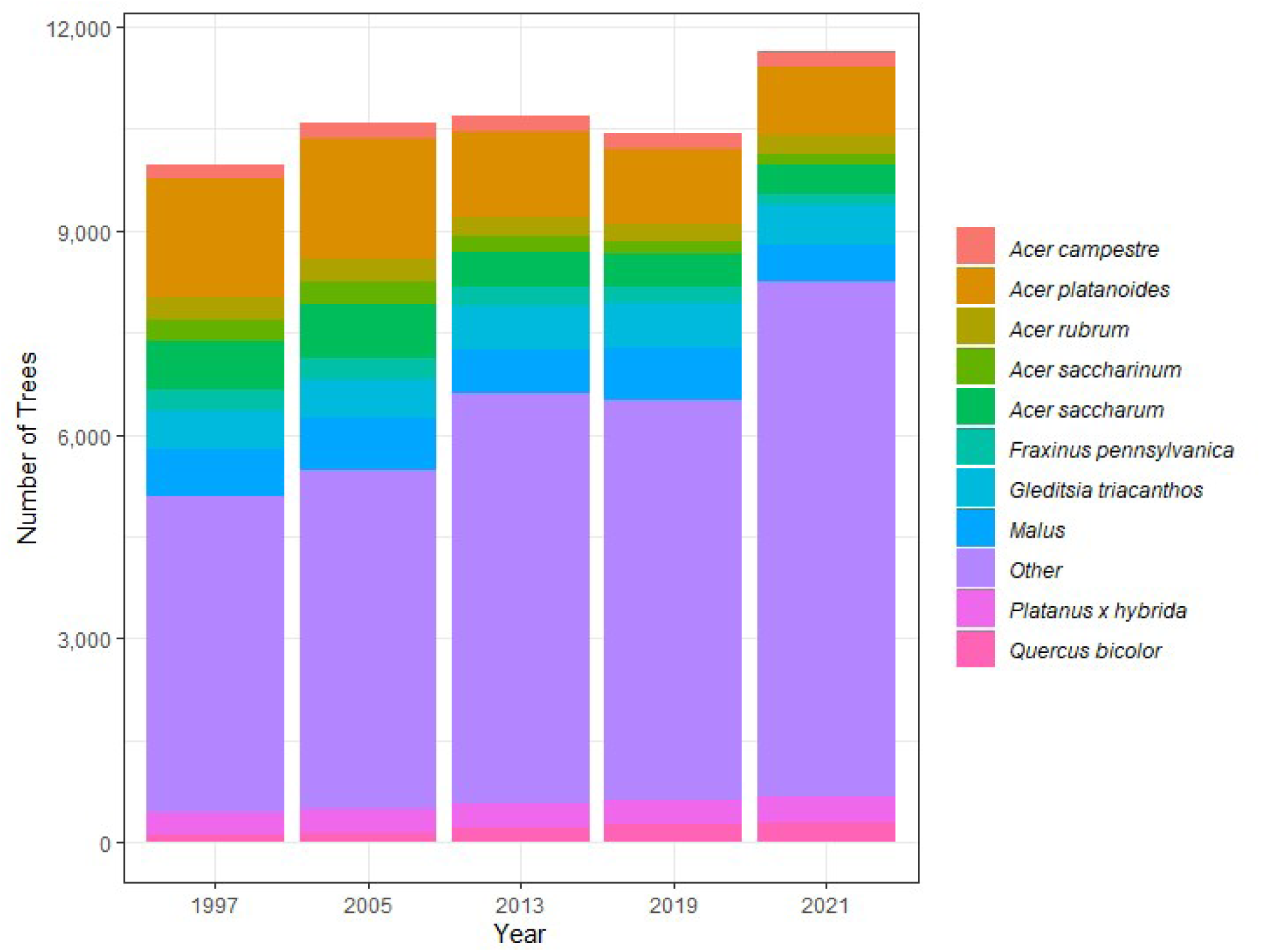
The total number of street trees in Ithaca, New York by species throughout the five most recent tree censuses. The 1928-1947 tree census was omitted as most of the trees recorded were identified only to genus.

**Supporting Information 3:**
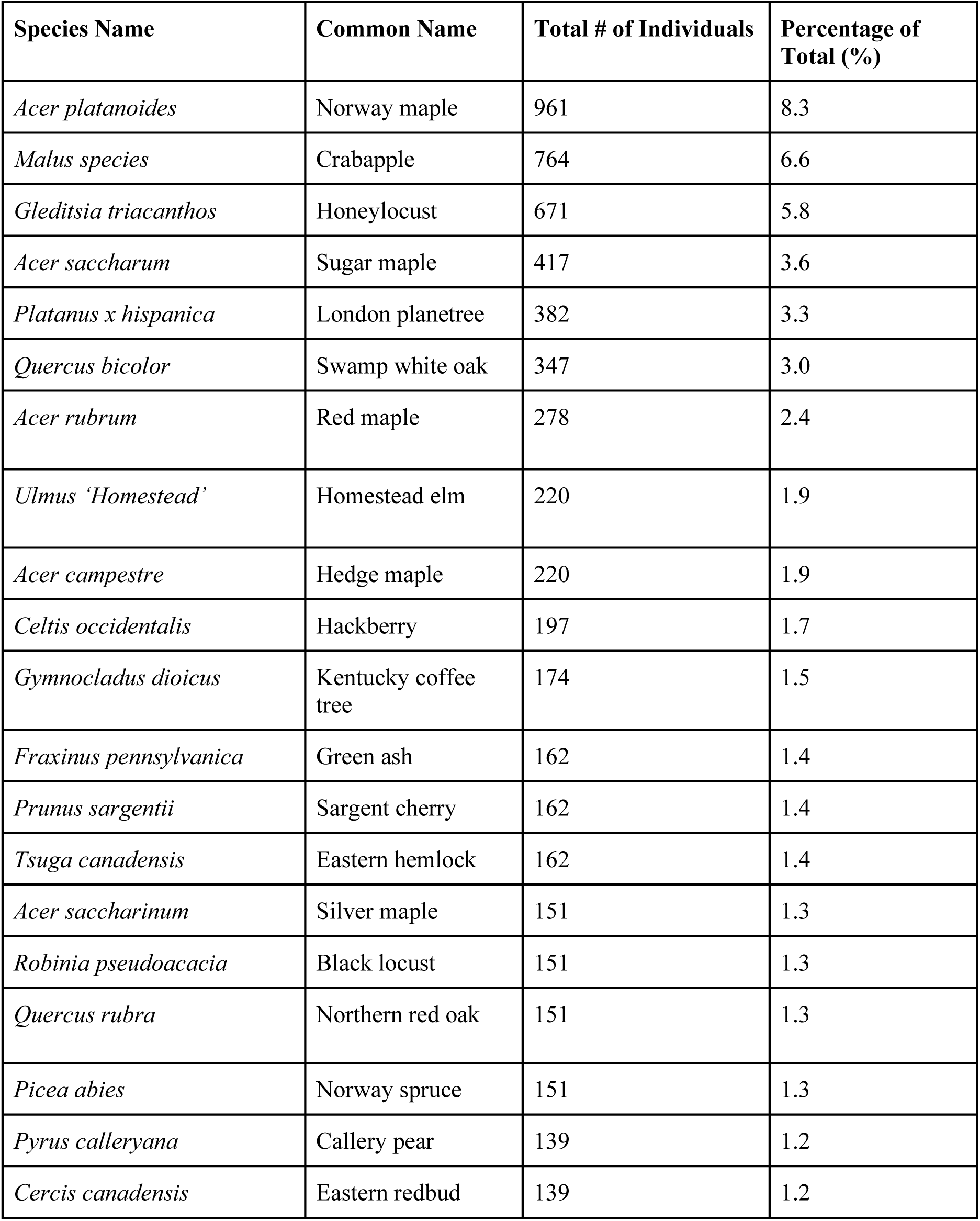
Table of species abundance from the most recent census (2021) for the most common tree species.

**Supporting Information 4:**
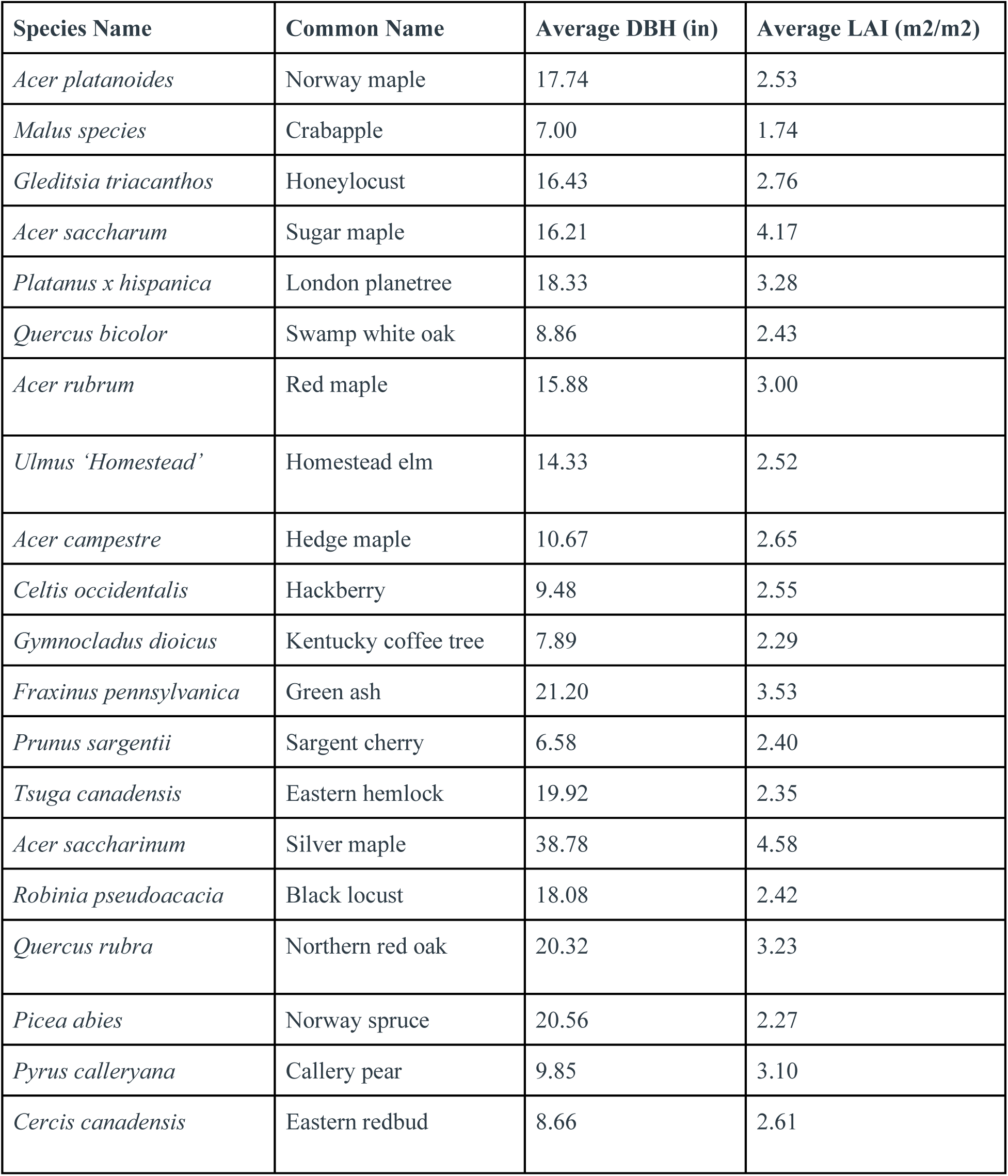
Average DBH and LAI for the most common tree species in Ithaca.

